# An abstract relational map emerges in the human medial prefrontal cortex with consolidation

**DOI:** 10.1101/2024.10.11.617652

**Authors:** Alon B. Baram, Hamed Nili, Ines Barreiros, Veronika Samborska, Timothy E. J. Behrens, Mona M. Garvert

## Abstract

Understanding the structure of a problem, such as the relationships between stimuli, supports fast learning and flexible reasoning. Rodent work has suggested that abstraction of structure away from sensory details occurs over the course of multiple days, in cortex. However, direct evidence of such explicit relational representations in humans is scarce, and it is unclear whether they emerge on similar timescales. Here, we combine a graph-learning paradigm with functional magnetic resonance imaging (fMRI) to look for such a relational map in the human brain. We first trained participants on two associative graphs with the same structure. We then scanned participants twice while they used this knowledge, with several days between scanning sessions. Using fMRI repetition suppression, we found an abstract relational representation in the medial prefrontal cortex (mPFC), that emerged across the two scanning sessions. This finding was also replicated using representational similarity analysis (RSA). These results shed new light on how neural representations organising relational knowledge change with time.

## Introduction

Humans and other animals constantly find themselves in situations they have never experienced before. How does our brain manage to choose adaptive behaviours given this diversity of experiences? Luckily, our world is replete with statistical structure^1^, which our brain represents as part of a “cognitive map”^2,3^. One particularly effective way of harnessing this structure for generalisation is to separate (or “factorise”) the representation of the environment’s relational structure from its specific sensory details^4^. A potential locus for this abstraction is the medial prefrontal cortex (mPFC), which has been implicated in recognising commonalities (“schemas”) across experiences^5–11^ and using these for inference^12^, as well as in concept learning^13,14^, structural generalisation of reinforcement learning problems^15–17^ and generalisation of spatial tasks across different paths^18–21^.

In addition, a recent line of research, inspired by rodent spatial studies, has shed new light on the relational properties of mPFC codes. Like the entorhinal cortex, the human mPFC contains grid cells^22^, which are active on a regular triangular lattice during spatial navigation^23^. Importantly, grid cells maintain their cell-to-cell activity covariance structure across different spatial environments^23–25^, suggesting they encode the statistical relationships common to all spatial environments. Notably, a flurry of studies using a technique to indirectly index grid cell activity suggests that the same brain network that displays “grid-like” activity during spatial navigation^26^ also displays it during navigation to novel “locations” in a wide range of non-spatial and abstract 2D tasks^27–31^. In most of these studies, the strongest grid-like signals emerge in the pregenual mPFC, suggesting this region encodes the 2D relational structure in these tasks. While these studies focused on 2D environments, theory suggests the 2D grid representation is merely a special case of abstract structural encoding applicable to a much wider range of task structures^4,32^. Whether the same cells that encode 2D structure (namely grid cells) also encode other structures is yet to be shown.

The mPFC has also been implicated in a related, yet distinct literature on abstraction via consolidation (i.e. on the time scale of days or weeks) across multiple memories^33,34^. For example, mPFC activity in rodents initially reflects specific perceptual features of learnt associations, and only later encodes generalisable relational features that are shared between associations^10,19,35^. However, the extent to which similar dynamics apply in humans remains unknown^36^, and a more precise understanding of the form of such abstract representations has remained elusive.

Here, we hypothesised that the mPFC abstracts an explicit relational representation of the environment over the course of multiple days. We used fMRI repetition suppression^37,38^ and representational similarity analysis (RSA)^39,40^ to assess whether such an abstract map emerges after training participants extensively on two graphs with different contexts, where the nodes are objects and the edges are associations between them. The objects and the graph structure were identical in both contexts, but pairwise distances between objects were decorrelated through their distribution on the two graphs. This allowed us to simultaneously test where the brain represents the task-relevant and irrelevant graphs in a task where relevance regularly switches. Crucially, we were also able to test for an abstracted graph representation which reflects the underlying common structure. To assess the role of consolidation in this abstraction, we examined these representations in two scanning sessions at least 24 hours apart.

We found that the medial temporal lobe represented the relevant graph in the first session and the irrelevant graph in both sessions, while a representation of the relational structure, independent of the specific stimulus identities, emerged in the mPFC in the second session. Such an abstracted map could presumably facilitate fast learning in new, but similarly structured situations.

## Results

### Subjects learn and flexibly switch between two graphs with the same structure

We designed a task to examine how the representation of abstract cognitive maps changes over the timescale of days. 23 participants were extensively trained on the relationships between a set of objects, each forming a graph (Figure 1A, B; training procedure outlined in the Methods section and Supplementary Figure S1). Importantly, the two graphs shared the same non-symmetric structure and differed only in terms of the distribution of objects across the graph’s nodes. Each object’s position on the blue graph thus corresponded to a unique object in the same position on the red graph. The objects were distributed such that pairwise distances between the objects were uncorrelated across the two graphs (Supplementary Figure S2A).

**Figure 1.**
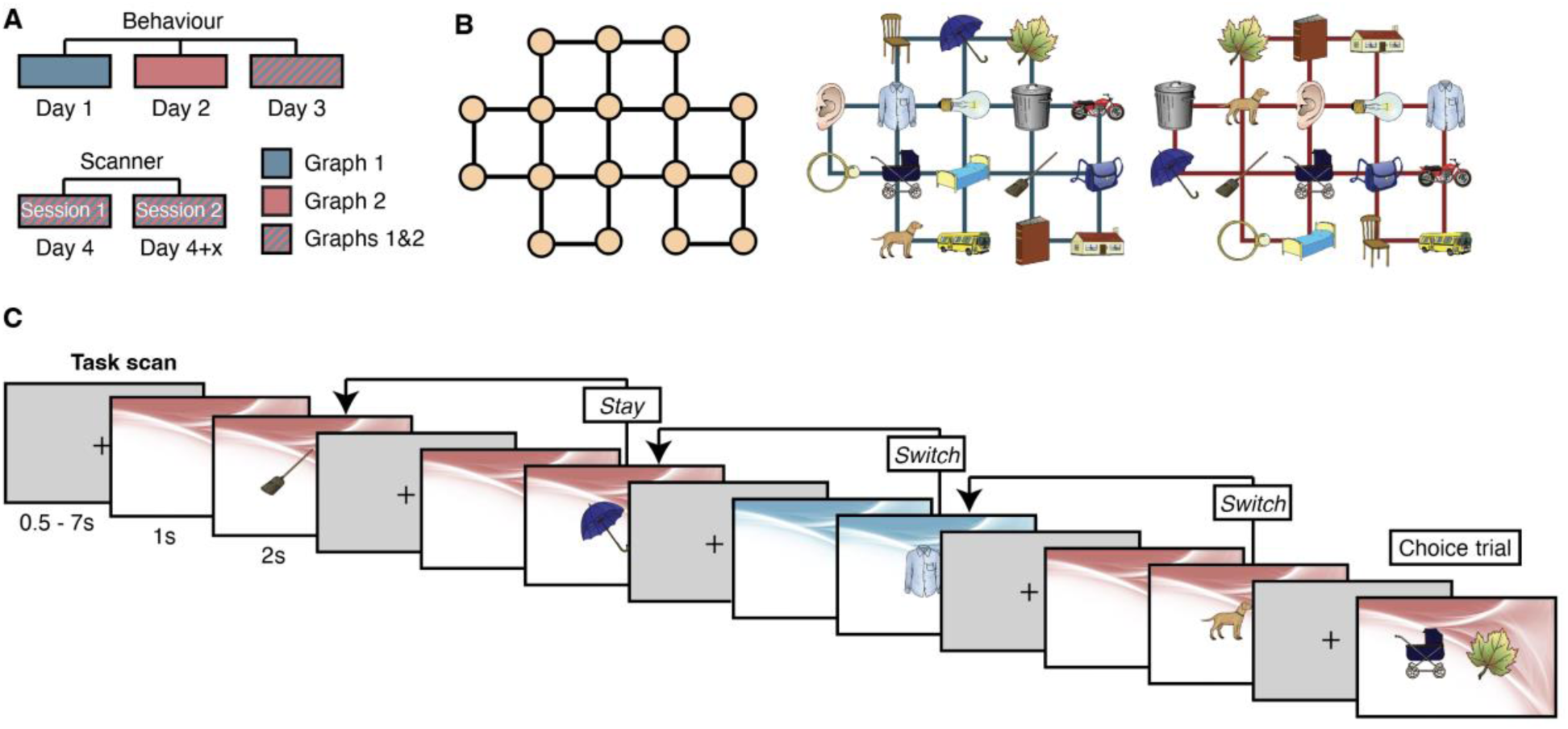
Experimental design. **A** Experimental design. Participants were trained online on the two graphs over three subsequent days (Supplementary Figure S1). On days 1 and 2, only one of the two graphs was trained, on day three both graphs were trained. Scan sessions took place on day 4 (session 1) and between one day and one week after the first session (session 2). **B** Graph structure (left) and example object distribution for the two graphs (middle and right). Across subjects, object distributions were randomised, and pairwise distances between objects on the two graphs were orthogonalised for every subject. **C** Task in the scanner. Participants were exposed to random sequences of objects after a brief context primer. Trials could be classified as *stay* trials (no change in context from trial to trial) and *switch* trials (change in context from one trial to the next). Occasionally, two objects were presented simultaneously. Participants were asked to indicate which of the two presented objects could be reached from the previously presented object in fewer steps in the relevant graph (*choice* trials).

Participants performed different tasks online, aimed at teaching them the pairwise associations between objects as well as sequences through the graph (Supplementary Figure S1). The underlying structure of the graphs was not made explicit. Each graph was learned separately on two subsequent days to minimise interference. Performance on the explicit structure-learning tasks did not differ between the two graphs learned on two separate training days (all p > 0.15, Supplementary Figure S1), suggesting that participants learned both graphs equally well. On the third day of training, participants repeated most of the tasks for both graphs to ensure that they learned to rapidly switch between them in a context-dependent manner. In all tasks, the context (i.e. the currently relevant graph) was indicated by a red or blue background image (order counterbalanced across subjects).

After three subsequent days of learning the object relationships, we presented subjects with sequences of objects in the scanner (Figure 1C). The first scanning session took place on the day after participants completed the training (session 1, day 4). The second scanning session took place between one day and one week after the first scanning session (session 2, day 4+x). The task in the scanner involved frequent context switches. The relevant context was always signaled by the same background color that subjects had learned to associate with a particular graph’s object distribution during the training procedure. On 12.5% of trials (once after each object), participants were asked to indicate by a button press which one of two presented objects could be reached in fewer steps from a preceding object on the currently relevant graph (*choice* trials). The purpose of this cover task was to keep participantś attention focused on the embedding of each presented object within its graph structure. In the object-difference task, sampled shortest-path distances spanned 1–6 steps in both the relevant and irrelevant graphs. Across subjects, mean sampled distances did not differ between relevant (M = 2.82, SD = 0.14) and irrelevant (M = 2.77, SD = 0.12) graphs (paired t-test, t_22_ = 1.23, p = 0.23). Thus, relevant and irrelevant distances were sampled equivalently in terms of range, mean, and variability.

Participants learned the graph structures very well, as demonstrated in a subset of participants (n = 19), who were asked to arrange both sets of objects relative to one another in an “arena”, after the last scanning session (Figure 2A-C). The pairwise distances in this arrangement correlated strongly with the ‘true’ link distances between pairs of objects (Figure 2D). Fisher-transformed correlations were significantly above zero for both graphs (Graph 1: median Fisher z = 0.74, IQR = 0.36–1.12; t_18_ = 6.64, 95% CI [0.49, 0.95], p = 0.000003, Cohen’s dz = 1.52; Graph 2: median Fisher z = 0.62, IQR = 0.48–1.05; t_18_ = 6.56, 95% CI [0.47, 0.92], p = 0.000004, Cohen’s dz = 1.51). Moreover, the correlation strengths between the two graph structures correlated with each other (r = 0.57, 95% CI [0.16, 0.83], p = 0.01, based on Fisher-transformed correlation coefficients) and were not significantly different from each other (difference in Fisher z: t_18_ = 0.25, 95% CI [-0.18, 0.23], p = 0.81, Cohen’s dz = 0.06). These results suggest that participants encoded both graph structures equally well. Indeed, we could recover both graph structures by applying multi-dimensional scaling to the distance matrix averaged across subjects (Figure 2B-C).

**Figure 2.**
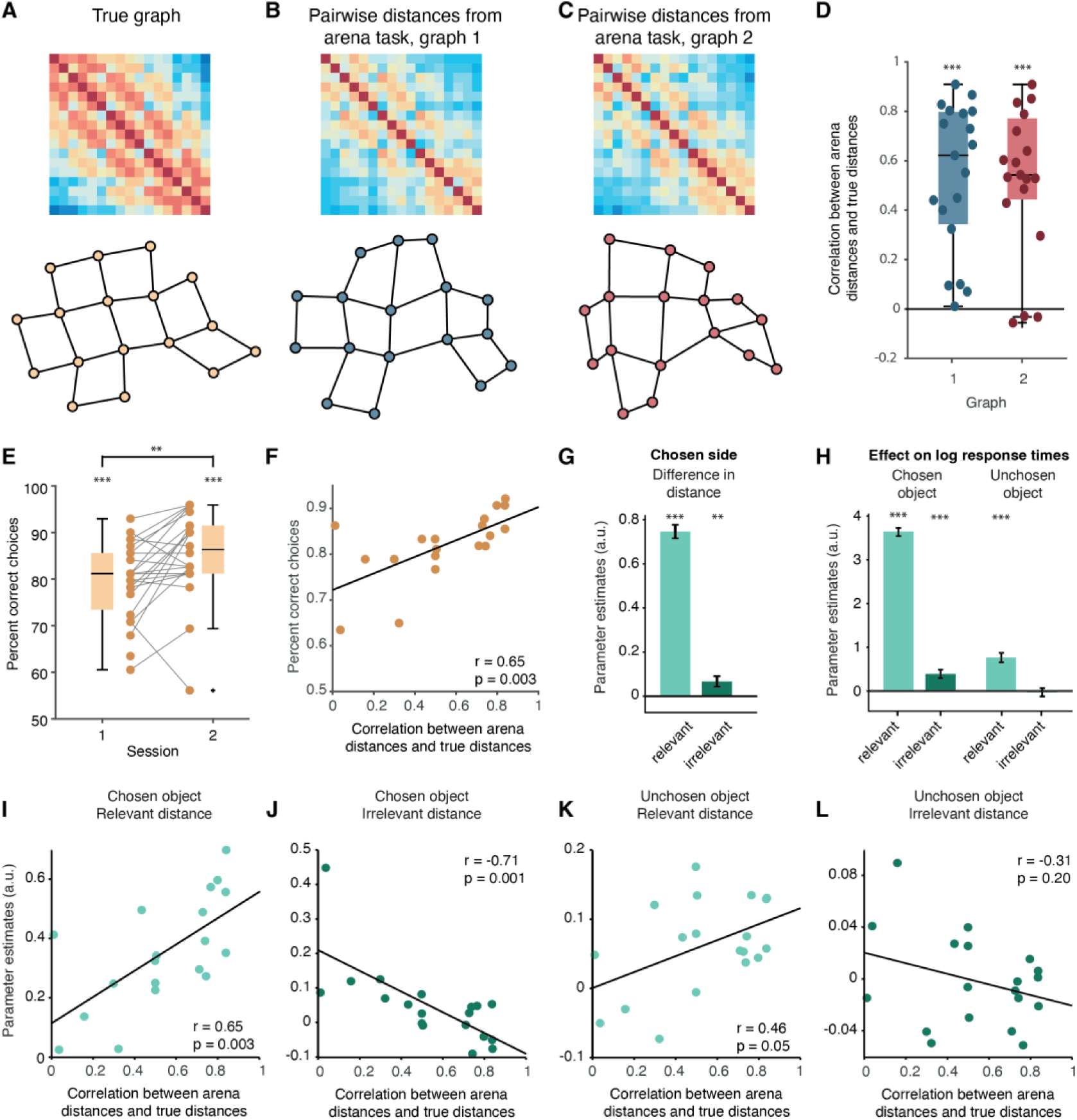
Behavioural performance. **A** Euclidian distances between object pairs (top) and reconstructed graph after projecting link distances into a two-dimensional space using multi-dimensional scaling (bottom). **B** Pairwise distances between objects of graph 1 as arranged by participants in an arena at the end of session 2, averaged across participants (top). Multi-dimensional scaling of the distance matrix was used to recover the graph (bottom). **C** Same as **B**, for graph 2. **D** Correlation between link distances in **A** and distances between positions in the arena for each graph (**B** and **C**). **E** Percent correct decisions in the choice task performed in the two scan sessions. Performance improves from session 1 to session 2. **F** Correlation between performance in the arena task and percent correct choices in the scanner, averaged across the two sessions. **G** Logistic mixed-effects regression coefficients reflecting the influence of the difference in distance to the chosen and unchosen objects on the relevant and irrelevant graph on chosen side. **H.** Linear mixed effects regression coefficients reflecting the influence of relevant and irrelevant shortest-path distances for chosen and unchosen objects on response times. In **G** and **H,** data from sessions 1 and 2 are pooled. **I** Correlation between arena performance and the influence of chosen/relevant distance on response times, averaged across the two sessions. **J** Correlation between arena performance and the influence of chosen/irrelevant distance on response times, averaged across the two sessions. **K** Correlation between arena performance and the influence of unchosen/relevant distance on response times, averaged across the two sessions. **L** Correlation between arena performance and the influence of unchosen/irrelevant distance on response times, averaged across the two sessions. Boxplots in **D** and **E** display the median and interquartile range as well as individual data points. *p < 0.05, **p < 0.01, ***p < 0.001

More importantly, participants also used their knowledge about the graph structures to solve the task in the scanner. Performance in the choice task was significantly above chance in both sessions (Session 1: t_22_ = 16.49, 95% CI [0.75, 0.83], p < 0.001, Cohen’s dz = 3.44; Session 2: Wilcoxon signed-rank W = 253, p < 0.001), with an improvement in performance from scanning session 1 to 2 (Figure 2E, t_21_ = 2.94, 95% CI [0.02, 0.09], p = 0.008, Cohen’s dz = 0.63). Across the two sessions performance in the choice task was highly correlated with performance in the post-scan arena task (Figure 2F, r = 0.65, 95% CI [0.28, 0.85], p = 0.003), suggesting that participants who had learned the graph structures particularly well benefitted the most from this knowledge in the scanner.

In a logistic mixed-effects model, with subject included as a random effect, the likelihood of choosing the left or right option was significantly influenced by the difference in distances between the target object and the objects on either side. The difference in distances on the relevant graph between the target object and the chosen object had a strong positive effect on the likelihood of choosing a given option (Figure 2G, β = 0.75, SE = 0.03, 95% CI [0.69, 0.81], t_3022_ = 24.12, p < 0.001). This suggests that participants were more likely to choose the option on the side that was closer to the target object on the relevant graph. A smaller, yet significant, effect was also observed for the difference in distances on the irrelevant graph between the target object and the two options (β = 0.07, SE = 0.02, 95% CI [0.02, 0.11], t_3022_ = 2.79, p = 0.005). This indicates that the irrelevant graph also had a measurable, though smaller, influence on decision-making compared to the relevant graph.

In a linear mixed-effects model with subject included as a random effect, the shortest-path distance between the target object and the chosen object on the relevant graph had a strong positive effect on log response time (Figure 2H, β = 0.36, SE = 0.01, 95% CI [0.34, 0.38], t_3020_ = 35.35, p < 0.001). Additionally, the shortest-path distance between the target object and the unchosen object on the relevant graph also had a significant positive effect on response time (β = 0.08, SE = 0.01, 95% CI [0.05, 0.10], t_3020_ = 6.87, p < 0.001). These results indicate that the length of mental simulation on the relevant graph, both from the chosen and unchosen options to the target, contributed significantly to longer response times. There was also a modest positive effect of the distance between the target object and the chosen object on the irrelevant graph (β = 0.04, SE = 0.01, 95% CI [0.02, 0.06], t_3020_ = 3.86, p < 0.001), suggesting some influence of irrelevant graph knowledge on response times. However, the distance between the target object and the unchosen object on the irrelevant graph did not significantly affect response times (β = −0.005, SE = 0.010, 95% CI [−0.024, 0.014], t_3020_ = −0.49, p = 0.62).

Next, we examined whether these effects of graph distances on response times in the scanner correlated with explicit knowledge of the graphs. We ran a separate regression for each participant (predicting, again, log response times with the chosen and unchosen distances on the relevant and irrelevant graphs) and correlated across subjects the resulting parameter estimates with an index of performance in the arena task - the correlation coefficients between the arena distances and “true” distances. We found that both the distance to the chosen object and the distance to the unchosen object on the relevant graph had a stronger influence on response times for participants who performed better in the arena task (chosen: r = 0.65, 95% CI [0.28, 0.85], p = 0.003; unchosen: r = 0.46, 95% CI [0.01, 0.76], p = 0.045, Figure 2I, K). This suggests that participants with a better understanding of the graph structures used this knowledge more effectively to solve the choice task.

Notably, the effect of distance to the chosen object on the irrelevant graph was negatively correlated with arena task performance (r = −0.71, 95% CI [−0.88, −0.38], p = 0.0006, Figure 2J), indicating that participants who knew the graph structures well were less distracted by irrelevant information. We found no significant correlation between performance in the arena task and the effect of unchosen distance on the irrelevant graph on response times, r = −0.31, 95% CI [−0.67, 0.17], p = 0.20, Figure 2L).

Together, these behavioural results suggest that participants had acquired and used structural knowledge about the relationships between objects to solve the cover task.

### Representations of both the relevant and irrelevant graphs in the MTL

To solve this context-dependent task, the brain must represent the object-embedded graph that is currently relevant in each trial. Multiple studies have shown that the medial temporal lobe (MTL) represents such associative graphs^41–43^ when these are task-relevant. Importantly, we and others have previously shown that MTL also represents such graphs even when learnt implicitly, and when subjects were engaged in a task for which the graph structure was irrelevant ^44,45^. We thus hypothesised that the MTL might simultaneously represent both the currently relevant and irrelevant graphs.

To test this, we used functional magnetic resonance imaging (fMRI) repetition suppression^38^, a technique based on the assumption that repeated stimuli evoke a suppressed response proportional to their distance in representational space^37^. To investigate within-graphs effects, we focused first on *stay* trials (where the context remains stable relative to the previous trial). We reasoned that in regions encoding a map-like representation of the relevant graph, the degree of fMRI suppression should decrease as a function of shortest-path distance on the relevant graph between the current and preceding object. (Figure 3A top). Meanwhile, in regions encoding the irrelevant graph, fMRI suppression *in the same trial* should scale with the shortest-path distance *on the other* (currently irrelevant) graph to the (same) preceding object (Figure 3A bottom). Since pairwise object distances were orthogonalised across graphs 1 and 2 (Supplementary Figure S2A), we were able to look at the representations of the relevant and irrelevant graphs simultaneously, in the same GLM.

**Figure 3.**
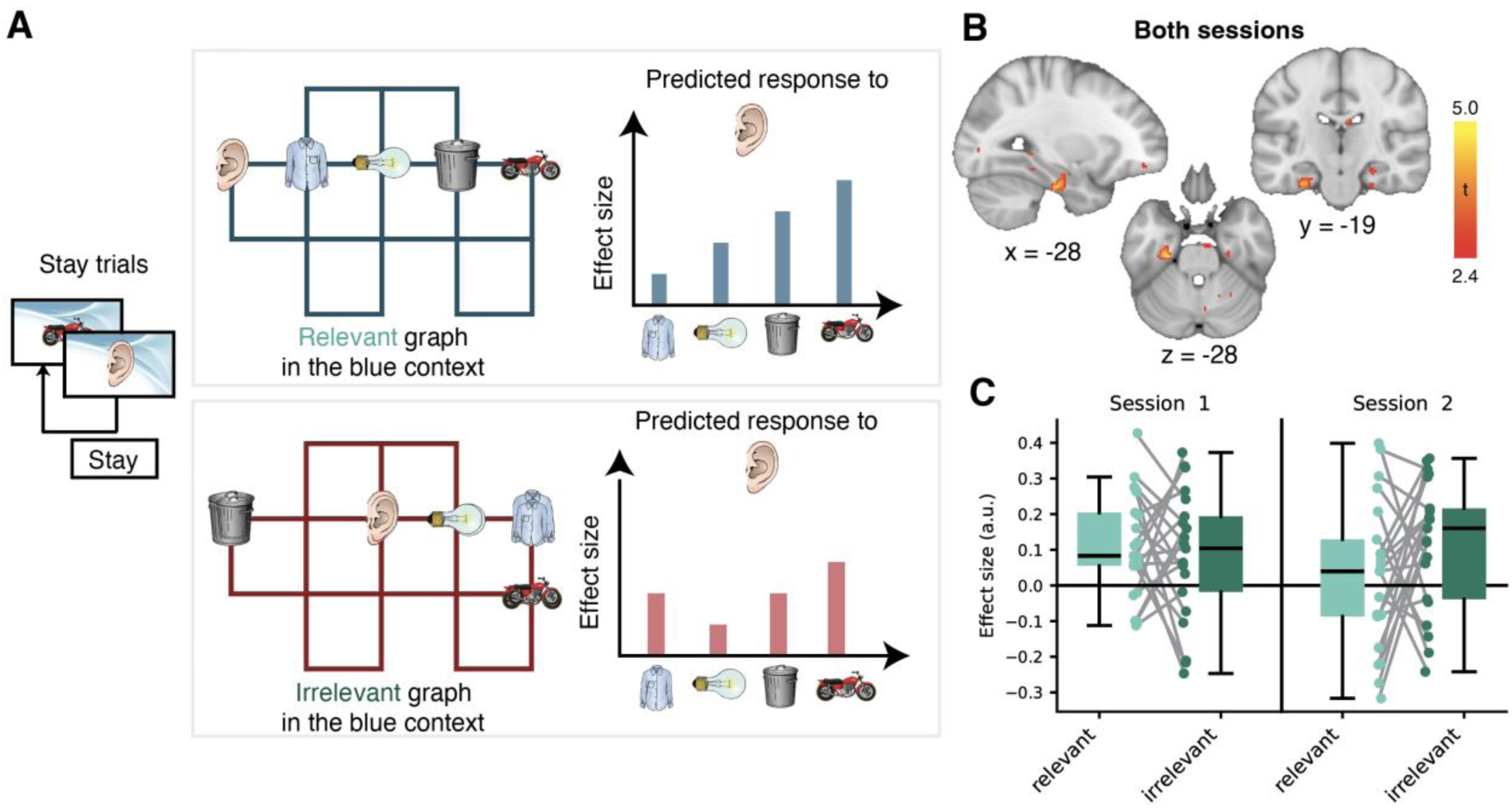
Relevant and irrelevant graph representations. A Analysis principle. On *stay* trials, distances between an object and its predecessor in the object sequence was defined on the relevant graph (here blue) and the irrelevant graph (here red). Visualised are distances from the Ear to four example objects. Since pairwise distances were orthogonalised for the two graphs, we were able to simultaneously assess the representation of relevant and irrelevant graphs. B MTL areas representing relevant and irrelevant distances across both sessions. C Visualisation of parameter estimates extracted from the left MTL cluster in B. Boxplots in C show the median, interquartile range, and individual participant values for each combination of Session and Graph relevance.

A contrast averaging both relevant and irrelevant distances across both sessions identified a cluster in the left MTL (Figure 3B-C). This cluster survived family wise error (FWE) small-volume correction (SVC) within a bilateral entorhinal cortex/subiculum mask (mask from ^44^, Supplementary Figure S2B, p = 0.03 corrected at peak level, peak t_22_ = 4.39, [−24, −22, −28]), and trended towards significance when SVC was performed in a larger MTL mask of bilateral hippocampus, parahippocampal cortex, and entorhinal cortex (p = 0.053 corrected at peak level, peak t_22_ = 4.94, [−26, −22, −28], mask in Supplementary Figure S2C). A cluster in the mPFC trended towards significance (Supplementary Figure S2F, p = 0.055 corrected at the cluster level in a mask of mPFC including Brodmann areas 14m and pregenual 32, mask in Supplementary Figure S2D, peak t_22_ =4.13 [−2, 38,−6]). In line with the comparable effects for graphs 1 and 2 across all our behavioural measures, we found no significant difference within a mask of the left MTL effect when comparing the suppression effects averaged across sessions for the two graphs (t_21_ = −1.41, 95% CI = [−0.06, 0.01], p = 0.17, Cohen’s dz = −0.30). A 2×2 ANOVA on the average parameter estimates in the effects’ clusters with the factors “Graph relevance” and “Session” did not reveal any significant main effects or interactions (Figure 3C, left MTL: Session: F_1,20_ = 0.60, p = 0.45; Relevance: F_1,20_ = 0.64, p = 0.43; Session × Relevance: F_1,20_ = 1.72, p = 0.21; Supplementary Figure S2F bottom, mPFC: Session: F_1,21_ = 1.99, p = 0.17; Relevance: F_1,21_ = 1.37, p = 0.26; Session × Relevance: F_1,21_ = 0.36, p = 0.55), suggesting that both the relevant and the irrelevant graphs were represented simultaneously in left MTL (and possibly mPFC), in both sessions. However, post-hoc tests suggest that the representation was absent in the second session for the relevant graph (one-sample t-tests vs 0: relevant, session 1: t_21_ = 3.99, p = 0.001; relevant, session 2: t_21_ = 0.09, p = 0.46; irrelevant, session 1: t_21_ = 2.31, p = 0.03; irrelevant, session 2: t_21_ = 2.85, p = 0.014; p-values Holm-Bonferroni corrected across the 4 tests; all pairwise two-sample t-tests: p > 0.06 uncorrected; p > 0.39 Holm-Bonferroni corrected across the 6 pairs of conditions; see Supplementary Figure S2E for the individual brain maps of each condition and further stats. Note that the one-sided tests against 0 have a selection bias which should inflate positive effects – and yet there is no [relevant, session 2] effect). In addition, we performed the same tests using average parameter estimates from bilateral masks of hippocampus and entorhinal cortex separately (see Methods for masks descriptions). These analyses did not reveal any significant differences within any single region, or between regions (p > 0.14 for all analyses).

Next, we wanted to corroborate this finding using an orthogonal analysis – representational similarity analysis (RSA^39^). Like repetition suppression, RSA also tests for the modulation of representations by hypothesised distances between conditions, but multivariately – using the correlation distances between voxel patterns. Importantly, repetition suppression and RSA are fully orthogonal. We used a mask of the left MTL cluster (thresholded at t=2.5, see inset in Supplementary Figure S2G-H) of the suppression analysis above as an ROI for RSA. We calculated correlations between voxel activation patterns (nVoxels = 103) of each pair of the 34 conditions (17 objects x 2 graphs), resulting in a 34 x 34 symmetric matrix (termed “Data Representational Dissimilarity Matrix”, or “Data RDM”, see illustration in Figure 4F for a similar construction process). To mirror the repetition suppression analysis, we focused only on the within-graph parts of this RDM (blue-to-blue and red-to-red), and constructed two hypothesis distance matrices (termed “Model RDMs”) from the relevant and irrelevant distances (e.g. distances on the blue and red graph, respectively, for the blue-to-blue part of the RDM, Supplementary Figure S2G, left). We then regressed the Data RDM onto the model RDMs. The contrast averaging both relevant and irrelevant distances across both sessions was significant in this ROI (one-sided t-test t_21_ = 2.318, p = 0.015; see further stats in Supplementary Figure S2G). Note however, that when this analysis was conducted without the visually confounded diagonal elements of the RDM, the effect only trended towards significance (one-sided t-test t_21_ = 1.514, p = 0.072, Supplementary Figure S2H).

**Figure 4.**
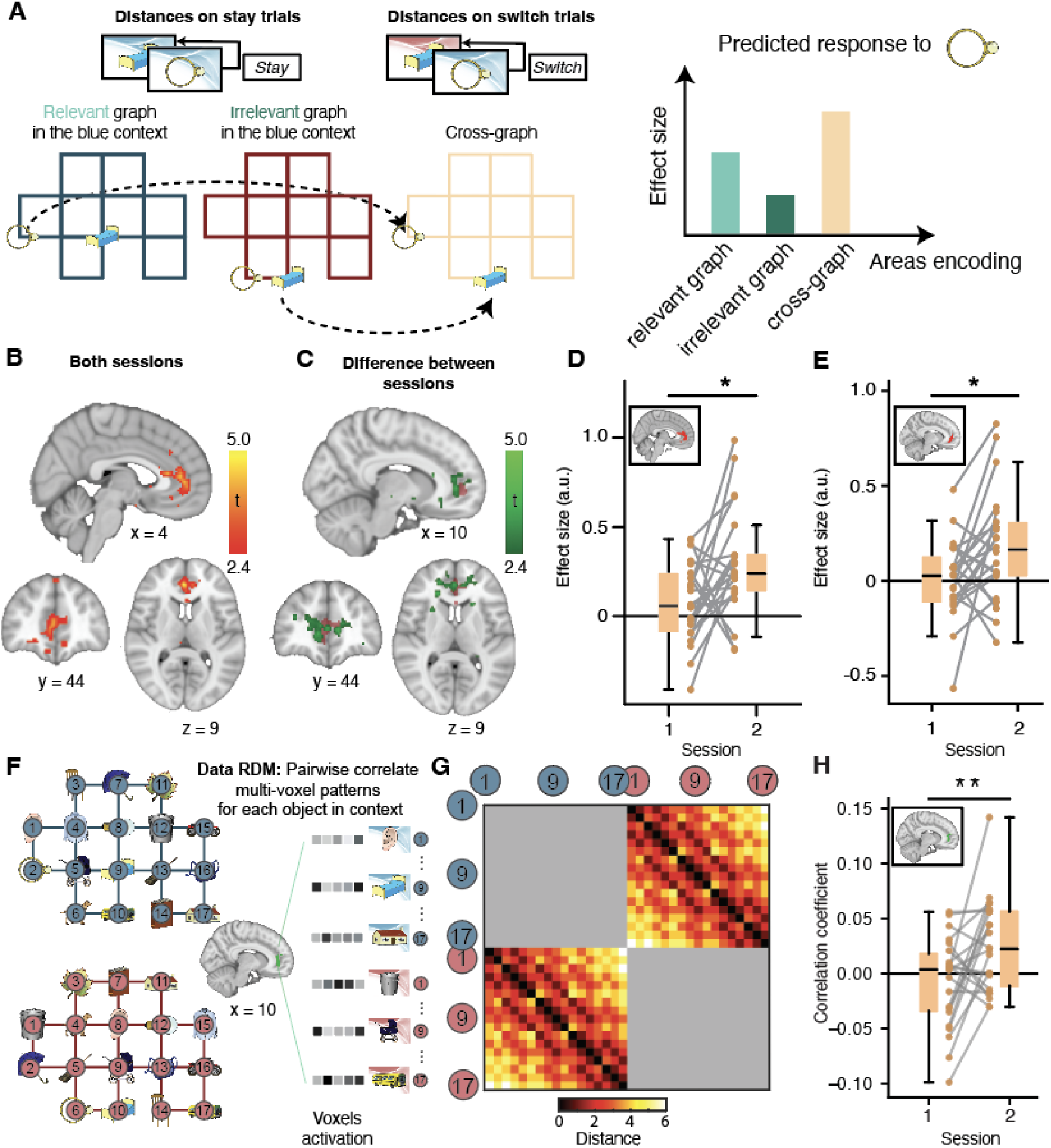
Abstract structure knowledge emerges in mPFC over time. **A** Logic of the cross-graph suppression analysis on *switch* trials. Cross-graph distances are defined as the distances between the position of the currently presented object on its graph and the position of the preceding object on the other graph. This measure corresponds to distances expected in an area that ignores the specific stimuli in a position, and instead reflects an abstract relational structure. **B** Across both sessions, cross-graph suppression is found in in the medial prefrontal cortex. **C** [session 2 > session 1] effect (green) overlaid on top of the mask extracted from **B** (red). **D** Visualisation of the effect from **C**: the cross-graph suppression effect is significantly higher in session 2 compared to session 1 in the ROI from the both-sessions effect (inset). **E** A cross-graph suppression effect also emerges in an ROI defined based on grid-like coding of abstract relationships (“bird space”)^29^. **F.** Representational Similarity Analysis (RSA) schematic. Using data from voxels in a mask of the thresholded [session 2 > session 1] cross-graph suppression effect (green effect in panel C), we extracted the dissimilarity of patterns across voxels separately for each of the 17 objects in each graph. **G.** Hypothesis matrix (Model RDM) of the cross-graph distances. Note that only conditions across graphs are compared. **H.** mPFC (see inset for mask) RSA effect for the analysis portrayed in panels F-G. Boxplots in **D, E, H** show the median, interquartile range, and individual participant values for the two sessions. *p < 0.05, **p < 0.01

### A generalisable abstract map in the mPFC

The medial prefrontal cortex (mPFC) is known to abstract the commonalities across experiences and represent schematic knowledge^5–7,17^. In addition, in humans mPFC contains grid cells^22^ and grid-like signals^26–31^ suggested to encode statistical structure in 2D spaces in a way that reflects commonalities and ignores particularities across tasks and environments^2,4^. Such abstract structure representations, grid-like or not, could be useful for transferring structural knowledge to new situations and thereby speed up learning.

In our experiment, such structure-coding would suggest the presence of an abstract map that is invariant to the specific context a stimulus was presented in. To test for such a cross-graph effects, we once again used repetition suppression, but this time on data from *switch* trials. We defined a cross-graph distance as the distance between the position of the currently presented object on its graph (Ring on Blue graph in Figure 4A) and the position of the preceding object on the other graph (Bed on Red graph). This distance reflects similarity in the abstract relational structure, regardless of stimulus identity. Such a cross-graph suppression effect would only be expected in brain areas that represent structure explicitly, and separately from the stimuli themselves.

A whole-brain analysis across both sessions identified a large significant cluster representing cross-graph distances in mPFC (Figure 4B, whole-brain FWE-corrected on the cluster level, p = 0.003, peak MNI [2, 42, 8] t_21_ = 5.1). This effect was driven by an emergence of a cross-graph suppression effect in session 2: within an ROI defined by the cross-graph effect across both sessions (threshold p < 0.01), the difference between sessions was significant (Figure 4C, [session 2 > session 1] SVC FWE-corrected on cluster level: p < 0.05, t_21_ = 3.66, peak MNI [4, 40, 12], t_21_ = 3.66; Figure 4D, t_21_ = 2.56, 95% CI = [0.02, 0.48], p = 0.02, Cohen’s dz = 0.55). This was also true when tested on the peak of the [both sessions] cross-graph effect (MNI [2, 42, 8], t_21_ = 3.1, p < 0.005). While these indirect repetition suppression effects do not allow us to make any inference about the particular types of cells involved in this representation, it is notable that these cross-graph suppression effects emerge across sessions in the same parts of mPFC showing grid-like coding of abstract relationships (mPFC ROI from Figure 2A of Constantinescu et al.^29^, [session 2 > session 1]: t_21_ = 2.38, 95% CI = [0.02, 0.32], p = 0.03, Cohen’s dz = 0.51, Figure 4E). Following a similar reasoning, it is worth noting that there was no emergent cross-graph effect when tested within an anatomical mask of the entorhinal cortex (one-sided t-test [session 2 - session 1]: p = 0.204, t_21_ = 0.84)).

Next, we wanted to test for the emergence of an abstracted map in mPFC using RSA (Kriegeskorte et al., 2008). We used a thresholded mask (t>2.5) of the [session 2 > session 1] cross-graph repetition suppression effect as an ROI (nVoxels = 264, Figure 4C green). As before, we calculated correlations between activation patterns in this ROI for each pair of the 34 conditions (Figure 4F). To assess whether a context-independent abstract map existed, we only used the across-graphs parts of the RDM (blue-to-red and red-to-blue), and ignored the within-graphs part (Figure 4G). We compared the data RDM of each scanning session to the Model RDM constructed from the cross-graph distances (illustrated in Figure 4A, data RDM in Figure 4G) using rank correlations. To mirror the repetition suppression results, we focused on the [session 2 > session 1] contrasts of the rank correlation coefficients. This contrast was significantly positive across participants (t_21_ = 2.87, p = 0.005, Figure 4H). There was no effect of this analysis when run in the left MTL ROI from Figure 3B (t_21_ = 1.12, p = 0.137). Taken together, these results provide converging evidence for an abstract relational map that emerges in mPFC between the two scanning sessions.

## Discussion

Here, we used a graph-learning paradigm to investigate how abstract relational structures are represented in the brain and how these representations change across two scanning sessions, days apart. Participants learned two different graphs characterised by the same underlying associative structures, but a different distribution of stimuli. The retrieval of the graphs – both currently relevant and irrelevant - was mediated by the MTL in the first scanning session, with the irrelevant graph retrieval also mediated by the MTL in the second session. In the mPFC, an abstract, stimulus-independent structural representation emerged in the days between the two scanning sessions. To our knowledge this is the first time the changes in such an explicit abstract map following consolidation have been investigated in humans. These findings were supported by evidence from both repetition suppression and RSA (though we note that the MTL RSA result, when controlled for visual confounds, was only trending towards significance).

Separating the representation of task structure from stimuli is useful, because it can facilitate the generalisation of knowledge across environments that share the same statistics^2,4,41^. If task knowledge is organised in such a factorised form, then solutions to new tasks do not have to be learnt afresh, they can instead be inferred^2,46^. Future research might directly test the relationship between the abstracted mPFC representation we found here and inference in novel situations sharing the same structure, for example by adding a test of behavioural generalisation on a novel, third graph (with the same structure) after the second scanning session.

We a-priori hypothesised that such an abstract representation might be found in the mPFC, as this region has been implicated in many human studies probing schematic and structural encoding^5,15,16,47^ as well as in the human cognitive maps literature^8,42^. In addition, together with the entorhinal cortex it is the most prominent area in which hexadirectional grid-like signals have been found in tasks with a 2D structure^26–31^, where a hexagonal grid is the most efficient form of structure representation^48^. With this in mind, it is noteworthy that we did not find an abstract representation in the entorhinal cortex. This might be due to differences in coding properties between the two regions, or simply due to the dropout of signal in the entorhinal cortex. We also emphasise that the relation between the abstract representation we find here and grid cells or grid-like coding is purely speculative – future work might explore this more directly, for example by measuring both representations in the same participants and testing for correlations between them.

Our results suggest that knowledge about the graph structure continued to be processed in mPFC after the first scan, even in the absence of further training. Moreover, it became more abstract in the second session, even though the task in the scanner only required knowledge of the specific graphs. This is in line with theories of systems consolidation positing that memories change qualitatively after acquisition, from an encoding of specific episodic information to the abstraction of statistical structure that is shared between episodes^35^. In this way, the neocortex gradually builds semantic knowledge of the world in a process that can take many weeks^34,49^. In rodents, such consolidation effects in the mPFC manifest themselves in a gradual loss of selectivity for perceptual features characterizing specific stimuli and an increase in the representation of relational features that are shared between stimuli^19^. In humans, featural overlap between memories was represented in mPFC (as well as hippocampus) one week after, but not immediately after encoding^36^. Importantly, we note that each participant’s first scan was performed when they were already extensively trained on the task, and that our sampling of time between scanning sessions was not continuous: each participant was scanned only twice, with anything between 1-6 days apart. Future research might address the time course of this emergence in further detail.

Indeed, memory reactivation during sleep plays a critical role in extracting rules and identifying abstract patterns across experiences^50,51^. A potential mechanism that might be involved in this abstraction - though not directly explored in this manuscript - is neural replay ^52^, which in humans reorganises experiences according to pre-learnt structures^53^. Replay also contains abstract structural codes in addition to sensory representations^53,54^, coincides with activation of the default mode network (including mPFC)^55^ and has been shown to be constrained by the structure of the task in a way that guides relational inference^56^. In addition, mPFC reactivation of category-level information during sleep in response to external sensory cues, supposedly triggering within-sleep replay ^57^, promotes memory consolidation^58^. Recent theoretical work suggests that the amount of replay of specific experiences depends on the reliability versus generalisability of the environment, thus balancing the trade-off between memory and generalisation^59^. Future work might explore a direct relationship between replay and the structural mPFC representations such as those we find here, for example by measuring replay in sleep between scanning days (using magnetoencephalogram and Temporally Delayed Linear Modelling^60^, and relating it to the strength of the emergent mPFC abstract representation.

While not the focus of the current manuscript, it is worth noting the rapid context-switching task subjects experienced in the scanner can be related to other proposed functions of the prefrontal cortex, such as cognitive control^61–63^. Indeed, in such tasks the ventromedial PFC contain rich representations of relevant and irrelevant contexts^64^. It is noteworthy that the strongest effects we found in this study – the repetition suppression cross-graph effects – were found when considering only in the *switch* trials, when the context changed. One possibility is that the abstract mPFC map is only needed and used in these more demanding trials. It is possible this map is still represented in *stay* trials, but in a weaker fashion. Weak support for this suggestion can be found in our trending repetition suppression effects of the relevant and irrelevant maps in the mPFC, particularly in session 2 (Supplementary Figure S2F).

In addition to the mPFC effects, we observed a reliable representation of both the relevant and the irrelevant graphs in the MTL, with the irrelevant map dominating the representation especially in session 2 (Figure 3B-C). Although one possible interpretation is that participants occasionally relied on the incorrect graph, the behavioural influences of the irrelevant graph are only modest in size. The dissociation between relatively strong MTL coding of irrelevant distances and weak behavioural effects suggests that neural representations in the MTL do not directly equate to behavioural influence. One possibility which is consistent with our previous findings^65–67^ is that the hippocampus maintains multiple relational structures in parallel, with task demands determining which representation is preferentially read out for behaviour^66^. Consistent with this account, an exploratory analysis revealed that in session 2, hippocampal coding of irrelevant distance was selectively enhanced on the first trial following a context switch (Supplementary Figure S3). This suggests a reconfiguration of relational representations during context updating, rather than a sustained use of the incorrect graph.

Another possible view on such context-switching tasks is through the lens of spatial cognition. In remapping experiments with rapid context switching (“teleporting”), the hippocampal representation rapidly flickers between the representations of both contexts, effectively implementing an inference process of the current context ^68^. While our manuscript does not focus on the hippocampus, we note that our MTL effect (partly including hippocampus) of the relevant and irrelevant graphs is compatible with such a remapping effect, as a representation of a relevant graph (which is strong particularly in session 1, see Figure 3C) is indeed what is expected from hippocampal remapping.

A key limitation of the present study is sample size. The final sample of 22 participants may limit sensitivity for detecting subtle representational effects, particularly in multivariate analyses and in smaller regions such as the MTL. At the same time, the central abstract-structure effects in mPFC survived whole-brain correction for multiple comparisons, indicating that they meet conservative statistical thresholds. In addition, the repeated-measures design was specifically suited to detecting within-participant change over time. Finally, we observed converging evidence for the same core conclusion using two independent analytic approaches (RSA and repetition suppression), reducing the likelihood that the findings reflect method-specific artifacts. Nevertheless, independent replication in a larger sample will be important to further establish the robustness and generalizability of these effects.

In conclusion, our results suggest that the medial temporal lobe represents both relevant and irrelevant graph relationships immediately after learning, while an abstract relational map that captures the underlying structure, independent of specific stimuli, emerges in the mPFC following several days. This provides evidence for the abstraction of structural knowledge in the human prefrontal cortex through consolidation.

## Resource availability

### Lead contact

Requests for further information and resources should be directed to and will be fulfilled by the lead contact, Alon B. Baram.

### Materials availability

This study did not generate new material contributions.

### Data availability

The subject-specific neuroimaging data reported in this study cannot be deposited in a public repository because consent for public data sharing was not obtained at the time of data collection, due to the ethics guidelines at the time (2017). Summary statistics describing these data, pseudonymised behavioural data and all original code have been deposited at https://github.com/alonbaram2/many-maps-alon. Any additional information required to reanalyse the data reported in this paper is available from one of the corresponding authors upon request.

## Acknowledgements

T.E.J.B. is supported by a Wellcome Principal Research Fellowship (219525/Z/19/Z), a Wellcome Collaborator award (214314/Z/18/Z), a JS McDonnell Foundation award (JSMF220020372), and by the Jean Francois and Marie-Laure de Clermont Tonerre Foundation. M.M.G. is supported by an ERC Starting Grant (101221509) the Interdisciplinary Center for Clinical Research (IZKF) at the University of Würzburg (Project number S-533). The Wellcome Centre for Integrative Neuroimaging and Wellcome Centre for Human Neuroimaging are each supported by core funding from the Wellcome Trust (203139/Z/16/Z, 203147/Z/16/Z). The Sainsbury-Wellcome centre is supported by core funding from the Wellcome Trust (219627/Z/19/Z) and the Gatsby Charitable Foundation (GAT3755).

## Author contributions

A.B.B., T.E.J.B. and M.M.G. conceived and designed the experiment and analyses. I.B., V.S. and M.M.G. collected the data. A.B.B. and M.M.G. conducted the analyses with support from H.N. A.B.B. and M.M.G. wrote the manuscript with input from T.E.J.B. T.E.J.B. acquired the funding.

## STAR Methods

### Key resources table

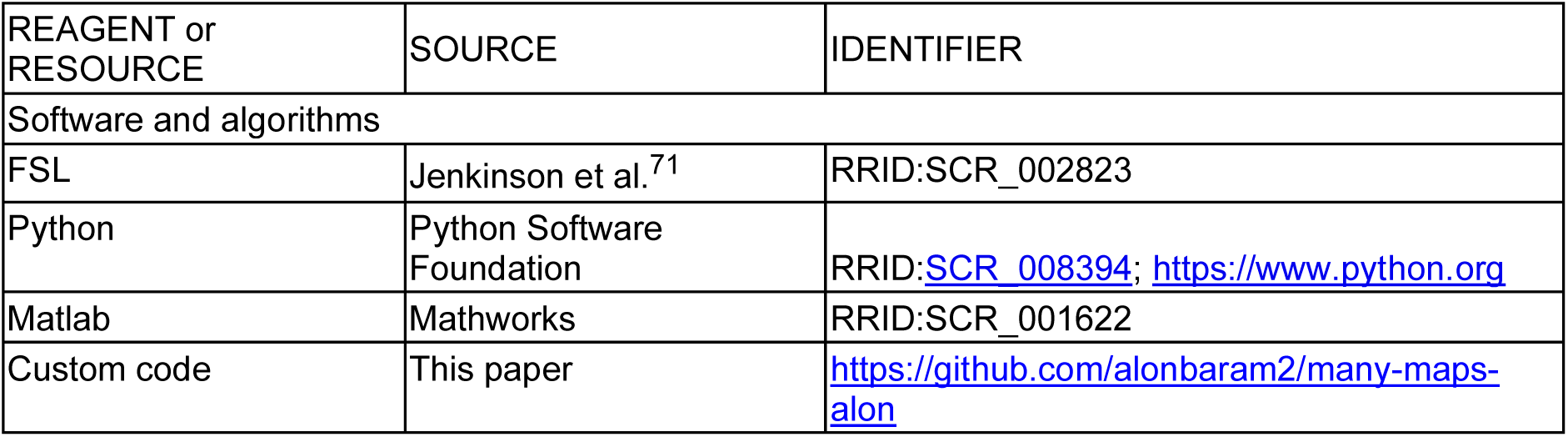

| REAGENT or RESOURCE | SOURCE | IDENTIFIER |
| --- | --- | --- |
| Software and algorithms |  |  |
| FSL | Jenkinson et al.<sup>71</sup> | RRID:SCR_002823 |
| Python | Python Software Foundation | RRID:SCR_008394; https://www.python.org |
| Matlab | Mathworks | RRID:SCR_001622 |
| Custom code | This paper | https://github.com/alonbaram2/many-maps-alon |

### Experimental model and study participant details

#### Participants

Twenty-three volunteers (aged 18 – 32 years, mean age ± standard deviation 23.2 ± 3.8 years, 14 female) participated in this study. One participant did not return to the lab for the second scan and only the first scan session of this participant was included in the analyses. All subjects reported normal or corrected-to-normal vision and no history of neurological or psychiatric disease. Participants gave written informed consent, and the study was approved by the Medical Sciences Inter-Divisional Research Ethics Committee (IDREC) of the University of Oxford (ref. number R49191/RE001). Participants were naïve to the purpose of the experiment. The scans took place at the Oxford Centre for Functional MRI of the Brain (FMRIB), Wellcome Centre for Integrative Neuroimaging.

### Method details

#### Experimental structure

The experiment took place over three training and two scan sessions (Figure 1A). The three training sessions and the first scan session took place over four subsequent days. The second scanning session was scheduled between one and seven days after the first scan, based on participant and scanner availability. In the training sessions participants performed a set of tasks to learn the relational structures. Participants were free to do the training in their own time at home through the online testing platform Prolific (www.prolific.com). These tasks were run in custom-written JavaScript, CSS and HTML scripts. Because of problems with data storage on Prolific, for some subjects a few task blocks during the pre-training could not be recorded. However, all participants completed all tasks.

#### Stimuli

Thirty-one coloured and shaded object images that were similar in terms of familiarity and complexity were selected from the ‘Snodgrass and Vanderwart ‘Like’ Objects’ picture set (http://wiki.cnbc.cmu.edu/Objects^69^. For each subject, a subset of 17 objects was randomly chosen and assigned to the 17 nodes of the two graph structures (Figure 1B). The positions of the objects on the two graphs were randomised such that pairwise distances between objects were orthogonal between graphs. The identity of the currently relevant graph was indicated by a blue or red background image. The colour associated with the graph learnt on the first day of training was randomly assigned and the alternative color was used for the second graph. Graphs one and two were initially learned separately on days one and two, and jointly on day three before the first scan.

#### Pre-training tasks

On each day, participants performed a set of computer-based tasks to learn the graph structures. In most of the tasks, participants could gain or lose coins depending on their performance. At the end of the experiment, the accumulated coins were converted to £ (every 100 coins = 1£ extra) and this money was paid in addition to the baseline payment of £7 per behavioural session. Visual feedback after each trial indicated whether a response was correct. On all days of the experiment, subjects were shown the number of correct trials and/or the number of coins collected at the end of each task.

To participate on subsequent days of the study, subjects were required to meet a minimum performance criterion of 400 coins collected on the first day of the experiment. This corresponds to about 70% of the total number of coins that could be obtained in the session. All trials and breaks between tasks were self-paced.

#### Tasks

The set of tasks was designed to teach participants the associations between pairs of objects and to familiarize them with the object transitions in the graph (Supplementary Figure S1A). Across tasks, the training procedure increased in complexity and abstraction: It progressed from learning local pairwise associations (Tasks 1,2 and 4) to exposure to transitions generated from the graph structure (Task 3) to multi-step inference and distance judgments (Tasks 5-6). By probing the same underlying graph structure using multiple different tasks, we aimed to promote robust encoding of the graph structure rather than task-specific strategies. The use of different tasks also minimized fatigue and ensured sustained attention across training. Participants were never instructed about the existence or the layout of the graph structure.

#### Task 1 - Object pair learning

Subjects were presented twice with each one of the 24 object pairs (i.e. objects directly connected to each other by a link in the graph structure) and asked to memorize each pair (Supplementary Figure S1B). This task taught pairwise associations without providing any information about the global organization of the objects. Participants were not informed that pairs formed a larger graph structure.

#### Task 2 - Associative memory task

Subjects were presented twice with each object pair as well as with 48 pairs of objects they did not learn to associate with each other in a random order and asked whether the two objects were paired or not (Supplementary Figure S1C). Subjects gained or lost a coin for each correct or incorrect response, respectively. This task reinforced accurate encoding of direct associations, without revealing higher-order graph structure.

#### Task 3 - Object orientation task

Subjects were exposed to object sequences generated from a random walk along the graph structure, such that each object would only be followed by a neighboring object on the graph structure ^44^. Objects were presented in one of two orientations, which were mirror-images of each other (Supplementary Figure S1D). Subjects performed an orthogonal orientation discrimination task, where they had to learn which orientation was associated with which button press. The random walk promoted integration of local associations into a broader relational representation. Subjects would gain or lose a coin for each correct or incorrect response, respectively. The task consisted of 100 trials.

#### Task 4 - Three-alternative forced choice task

Subjects were presented with one object at the top of the screen and had to pick the object it was paired with from three objects presented below (Supplementary Figure S1E). Each object was presented as many times as the number of objects it was linked to on the graph, such that each association was tested once. Subjects would gain or lose a coin for each correct or incorrect response, respectively. Like the associative memory task, this task reinforced direct connections between objects to strengthen edge knowledge.

#### Task 5 - Directed navigation task

In this task subjects navigated through the graph structure to find a given target object. The target object was displayed at the top of the screen. In the center of the screen the object corresponding to the current state were presented. Above this object, two cards were presented which were paired with this object. When choosing the next object, the object at the center was replaced by the selected object and the options were replaced by two new objects the chosen object was paired with (Supplementary Figure S1F). This means that by clicking through the objects, subjects moved from association to association until they reached the target. Below the objects, a progress bar and a number of coins were presented. At the start of a trial, the number of coins corresponded to the distance on the graph between the starting and the target object. Selecting the object option closer to the target would move the bar by the distance corresponding to (the total length of the bar/minimum number of steps to reach target) to the right. Conversely, incorrect choices would move the progress bar to the left and a coin was subtracted from the amount that could be earnt on the trial. When subjects filled the progress bar, they got the number of coins displayed below. This means that the fewer steps it took a subject to reach the target, the more coins they gained. At the end of each trial, subjects were presented with the sequence of objects they had chosen to encourage learning. This task consisted of 30 trials. This task required participants to integrate local pairwise knowledge across multiple steps and enabled us to probe whether participants had formed a global representation of the graph structure. Unlike all earlier tasks, successful performance required multi-step relational inference.

#### Task 6 - Distance difference task

On each trial, participants were presented with a target object at the bottom of the screen and two goal objects presented on top of the target object. Participants were required to select the option that they would reach in fewer steps if they moved from association to association like in the directed navigation task from the target object (Supplementary Figure S1G). In other words, participants were required to select the option closer to the target object on the graph. Participants would gain or lose a coin for each correct or incorrect response, respectively. This task consisted of 30 trials. This task assessed whether participants represent relational distance or only direct connections.

#### fMRI experiment

Before the scanning session on day 4, participants performed a brief reminder training session consisting of one block of task 4 (three-alternative forced choice task) and one block of task 6 (goal navigation task). They were also familiarized with the task they would perform in the scanner through a brief pre-scan training session that was the same as the task performed in the scanner (15 trials).

In the scanner, subjects were presented with random object sequences. Each of the 17 objects was presented exactly four times per graph in each experimental block, resulting in a total number of 17 x 2 x 4 = 136 presented objects per block. There were 4 experimental blocks. 1 second before stimulus onset, the background colour changed according to the relevant graph. Stimuli were then presented with the stimulus in context for 2 seconds, followed by a jittered inter-trial interval (ITI) generated from an exponential distribution with a mean of 1.7 seconds and truncated between 0.5 and 7.5 seconds (Figure 1C).

While observing the object sequences subjects performed a cover task of infrequently reporting by button press which one of two presented objects they would reach in fewer steps. This choice task occurred after random intervals exactly once after each object, i.e. in 12.5% of the total number of trials. Each correct button press was rewarded with £0.10 paid out in addition to the baseline fee to ensure that subjects attended to the stimuli. Subjects received a brief training on this task before they performed it in the scanner.

#### Arena task

At the end of the second scanning session, a subset of all participants (n=19) performed the Arena task separately for each context^70^. Here, all stimuli were initially displayed in random order outside a circular arena. Participants were instructed to arrange the objects inside the arena such that objects that were close to each other in the underlying graph structure were placed close together spatially. The item sets presented on each trial were adaptively selected using a “lift-the-weakest” heuristic, which optimizes subset construction based on previously collected judgments^70^. The resulting dissimilarity matrix, representing each participant’s perceived relationships between all items, was estimated by combining repeated judgments across pairs.

#### fMRI data acquisition and pre-processing

Visual stimuli were projected onto a screen via a computer monitor. Subjects indicated their choice using an MRI-compatible button box. MRI data were acquired using a 32-channel head coil on a 3 Tesla Magnetom Prisma scanner (Siemens, Erlangen, Germany). A T2*-weighted echo-planar imaging (EPI) sequence with a multi-band acceleration factor of 3 and within-plane acceleration factor (iPAT) of 2 was used to collect 72 transverse slices (interleaved order) with TR = 1.235 s, TE = 20 ms, flip angle = 65°, voxel resolution = 2×2×2 mm. Slices were tilted by 30° relative to the axial axis. The first five volumes of each block were discarded to allow for scanner equilibration. After two fMRI blocks, a T1-weighted anatomical scan with 1×1×1 mm resolution was acquired. In addition, a whole-brain field map with dual echo-time images (TE1 = 4.92 ms, TE2 = 7.38 ms, resolution 3×3×3 mm) was obtained to measure and later correct for geometric distortions due to susceptibility-induced field inhomogeneities. We performed slice time correction, corrected for signal bias, and realigned functional scans to the first volume in the sequence using a six-parameter rigid body transformation to correct for motion. Contrast images were later spatially normalized by warping subject-specific images to an MNI (Montreal Neurological Institute) reference brain and smoothed using a 6-mm full-width at half maximum Gaussian kernel. All preprocessing steps were performed with FSL^71^. Note that for the RSA analyses we used unsmoothed preprocessed data and later smoothed the resulting data RDMs (see below).

### Quantification and statistical analysis

#### Behavioural data analysis

All data analysis of the training sessions, as well as of performance in the arena task after session 2, was performed using custom-written Matlab (MATLAB R2022b, Mathworks, USA) or Python scripts (Python 3.11.4).

To obtain a metric for performance in the arena task, we first organized each subject’s responses into pairwise distance matrices for each pair of objects. For visualisation purposes, we projected the matrices to a two-dimensional space using multi-dimensional scaling (MDS, Figure 2A-C bottom). Next, we correlated these distances with the “true” distance matrix, based on the number of shortest paths between object pairs (Figure 2D), as well as correlating the matrices of the two graphs with each other. To test for significance of these correlations across subjects we ran a two-sided t-test on the Fisher-transformed correlation coefficients.

Analysis of the behaviour in the choice trials during the scans was performed using MATLAB R2022b. We fitted a logistic mixed-effects model using the fitglme function, with subject included as a random effect. The dependent variable was the binary choice (left vs. right option), and the independent variables were the differences in distance between the target object and the objects presented on each side (Figure 2G).

We then ran a linear mixed-effects model in MATLAB using the fitlme function, with subject included as a random effect. The dependent variable was the response times in the choice trials and the four independent variables were the distances between the target object and the 1) chosen object on the relevant graph; 2) unchosen object on the relevant graph; 3) chosen object on the irrelevant graph; 4) unchosen object on the irrelevant graph (Figure 2H). Next, we wanted to obtain subject-by-subject scores of the effects from this analysis, in order test for a correlation with the performance in the arena task (Figure 2I-L). We achieved this by running a separate linear regression for each subject, with the same dependent and independent variables as above.

All statistical tests were two-sided, unless mentioned otherwise. Repeated measurements from the same participants were modelled using paired tests or mixed-effects models with subject as a random effect. Normality of the relevant distributions was assessed using the Shapiro–Wilk test; parametric tests (t-tests) were used when normality was not rejected, and non-parametric alternatives (Wilcoxon signed-rank tests) were used otherwise.

#### fMRI data analysis: Repetition suppression

We implemented an event-related GLM in SPM 12 to analyse the fMRI data. We modeled the onsets of objects separately for each graph and for *stay* and *switch* trials. Each regressor additionally contained two parametric modulators, one reflecting shortest-path distance between the current object and the preceding object on the relevant graph and one reflecting the shortest-path distance between the current object and the preceding object on the irrelevant graph. These two regressor were designed to be uncorrelated (though in practice had small correlations, see Supplementary Figure S2A). Serial orthogonalisation in SPM was turned off for all GLMs. Contrasts were defined as [relevant] + [irrelevant] distances on *stay* trials (Figure 3) or as [relevant] distances on *switch* trials (Figure 4). The contrast images of all subjects from the first level were analysed as a second-level random effects analysis.

#### Representations of the relevant and irrelevant graphs

We report our results in the hippocampal–entorhinal formation and medial prefrontal cortex, as these were our a priori ROIs for the representations of the relevant and the irrelevant graphs, at a cluster-defining statistical threshold of p<0.01 uncorrected, combined with SVC for multiple comparisons (peak-level family-wise error [FWE] corrected at p<0.05). For the SVC procedure, we used three different anatomical masks. The first mask consisted of the entorhinal cortex and subiculum alone and was received with thanks from Chadwick et al. ^72^, Supplemental Figure S2B). The second mask also contained other medial temporal lobe regions implicated in encoding physical space and comprised the hippocampus, entorhinal cortex, and parahippocampal cortex, as defined according to the maximum probability tissue labels provided by Neuromorphometrics, Inc. (Supplemental Figure S2C). The third mask was an mPFC mask obtained by the union of the masks of Brodmann areas 32sg and 14m from the Brainnetome atlas^73^, thresholded at 50 (Supplementary Figure S2D). Activations in other brain regions were only considered significant at a cluster-defining threshold of p<0.01 uncorrected if they survived whole-brain FWE correction at the cluster level (p<0.05). While no areas survived this stringent correction for multiple comparisons, other regions are shown in Figures 3B, 4B and 4C at an uncorrected threshold of p<0.01 for completeness. While we used masks to correct for multiple comparisons in our ROI, all statistical parametric maps presented in the manuscript are unmasked. All the clusters used in the manuscripts for extraction of (independent) parameter estimates were obtained by thresholding at t=2.5. To test whether suppression effects differed between the two graph structures, we extracted four condition estimates from the fMRI first-level models: graph 1 relevant, graph 2 relevant, graph 1 irrelevant, and graph 2 irrelevant. For each participant and session, the mean beta value within the MTL mask was obtained. Outliers exceeding ±3 SD within each session × condition were excluded. To ensure balanced data, only participants contributing all four conditions were retained. We tested for session- and relevance-specific effects using a 2×2 repeated measures ANOVA. For each participant, we then averaged the relevant and irrelevant conditions within each graph to obtain a single graph 1 and graph 2 suppression estimate. The resulting paired values were compared using a two-tailed paired t-test, testing the null hypothesis of no difference between the two graphs. We additionally performed post-hoc analyses to test for effects of each of the 4 conditions (relevant/irrelevant, session 1/session 2, four one-sided t-tests against 0, corrected for multiple comparisons using Holm-Bonferroni correction) as well as differences between each pair of conditions (six two-sided paired t-tests, corrected for multiple comparisons using Holm-Bonferroni correction). Finally, we ran both the ANOVA analysis and the post-hoc analyses using the mean parameter estimates, as well as the within-subject difference of means, of two additional anatomical masks: bilateral entorhinal cortex (From the Juelich Histological atlas, thresholded at 25), and bilateral hippocampus (Harvard-Oxford atlas, thresholded at 25).

#### fMRI data analysis: Representational Similarity Analysis

Representational similarity analysis^39^ was performed in ROIs derived from repetition suppression effects. These analyses were constructed to ask the same questions as their corresponding repetition suppression analysis, but with an independent measure. We transformed the preprocessed fMRI data to MNI space, and ran a GLM where we modelled the presentation of each object in each of the graphs as a separate condition (without any distinction between *stay* or *switch* trials). Note that here, unlike in the repetition suppression analysis, we did not average the betas for each condition across experimental runs, as we will use these beta maps to do RSA across runs. Representational dissimilarity was computed as the correlation distances between voxel patterns in an ROI after spatially whitening the data within the ROI^74^. To account for temporal autocorrelations and biases due to temporal proximity, we correlated data across experimental runs and averaged the correlation coefficients to obtain a single 34 x 34 conditions RDM per subject per session. This procedure was performed twice: for the ROI from the left MTL (lMTL) cluster of the relevant + irrelevant distances effect at stay trials across both scanning sessions (from Figure 3B, thresholded at t > 2.5, number of voxels in mask = 103); and the mPFC [session 2 > session 1] cross-graph effect at switch trials (green effect in Figure 4C thresholded at t > 2.5, number of voxels in mask = 264).

Next, we compared the data RDM in the ROI to the hypothesis model RDMs. Using linear regression, we compared the MTL data RDM with model RDMs constructed from the relevant and irrelevant distances within each graph (cross-graph RDM elements were excluded). The analysis was performed either including the diagonal (Figure S2G, left) or excluding it (Figure S2H). We then used the betas from this regression in a similar way as we did for the (averaged across ROI) repetition suppression effect. We first tested for significance of the [relevant and irrelevant distances, across both sessions] effect by averaging the betas across the four conditions and performing a one-sided t-test against 0. Next, we tested for session- and relevance-specific effects, as well as an interaction, using a 2×2 repeated measures ANOVA with factors “session” and “relevance”.

For the mPFC [session 2 > session 1] cross-graph effect, we used a similar procedure. However, because this analysis involved only a single model RDM, (relevant distance across graphs, Figure 4G) we used Kendall’s Tau rank correlation rather than a regression. We then performed a one-sided paired t-test on the tau coefficients against 0, testing the hypothesis that the data RDM is more correlated with the model RDM in session 2 than in session 1.

