## Supplementary Figures for "An abstract relational map emerges in the human medial prefrontal cortex with consolidation"

### Supplementary Figure 1

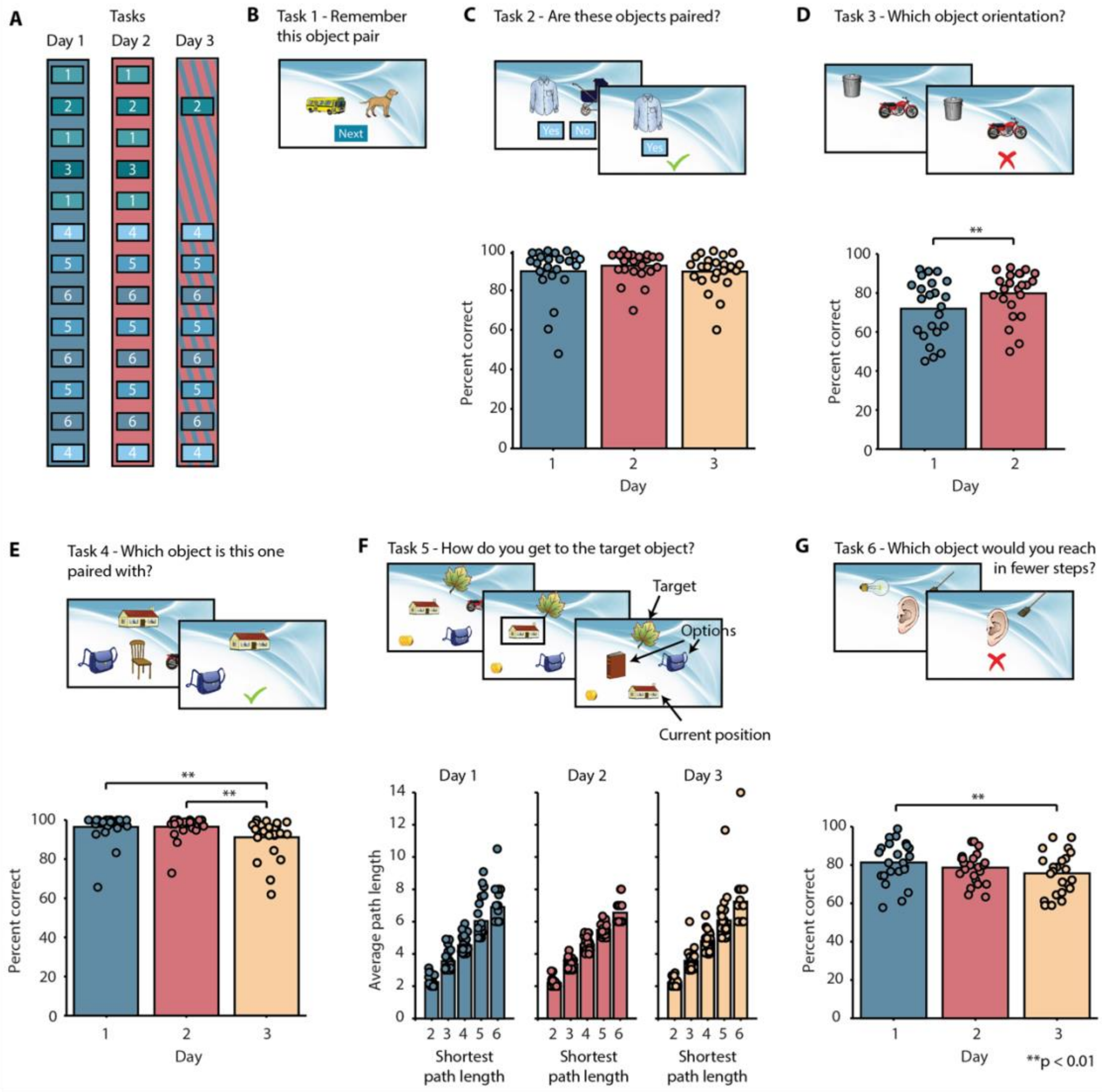


**Figure S1. Behavioural training procedure and explicit knowledge of the graph structures. A** Participants completed a sequence of training tasks across three consecutive days designed to promote and probe relational knowledge of two graph structures. Importantly, participants were not instructed about the existence or topology of the graphs; knowledge of the relational structure had to be derived from object associations and graph-constrained transitions. Training progressed from reinforcement of local pairwise associations (B,C,E) to tasks requiring multi-step relational inference and distance comparison (F–G). No time limit was imposed on any of the tasks and feedback was displayed after every trial. On day 1, graph 1 was used as a basis for generating pairs and sequences in **B-G**, on day 2, graph 2 was used and on day 3 trials of the two types alternated. **B** Task 1. Participants were presented with two objects corresponding to objects connected by a link on the graph and instructed to memorise the object pair. Each object pair was assessed once per block. **C** Task 2. Participants were asked whether two objects were paired or not. Each object pair was assessed once per block. Participants performed significantly above chance on all three days ($p < 0.0001$ for all days). A repeated measures ANOVA with a Greenhouse-Geisser correction determined that performance did not differ statistically between days ($F_{1.334,\ 29.349} = 1.63$, $P = 0.22$). **D** Task 3. Participants were exposed to object sequences generated from a random walk on the graph (in the example shown, the bin was the previous object in the sequence and the motorbike is the current object). While performing an orthogonal orientation discrimination task, they experienced transitions that reflected the underlying graph structure. This design promoted integration of local associations through structured exposure without explicitly describing the graph layout. Participants performed significantly better than chance on both days (both $p < 0.001$) and improved from day 1 to day 2 ($t_{22} = 2.9$, $p = 0.008$). **E** Task 4. Three-alternative forced choice task. Participants had to identify an associated object from a set of three. Each object pair was tested once per block. Participants performed significantly above chance on all three days ($p < 0.001$ for all days). A repeated measures ANOVA with a Greenhouse-Geisser correction determined that performance differed statistically significantly between days ($F_{1.553,\ 34.160} = 4.78$, $P = 0.02$), because performance on day 3 was worse than on days 1 and 2 (both $p = 0.003$, post hoc tests using Bonferroni correction). **F** Task 5. Participants were instructed to click from a start object displayed at the bottom to a target object displayed at the top in as few steps as possible by repeatedly choosing one of two objects presented in the centre. The chosen object would become the next start object. Every time an incorrect option was chosen, the progress bar would be reduced in size. At the end, an amount of coins corresponding to the length of the progress bar was offered to the participant. Bottom: the average number of steps taken, split according to the shortest possible path length for each training session. **G** Task 6. Participants were requested to mentally simulate a path through the graph structure and choose one of two objects displayed at the top that could be reached in fewer steps from the starting object displayed at the bottom. This task required explicit comparison of shortest-path distances, assessing whether participants encoded graded relational distance beyond direct associations. This task was later also performed in the scanner. Participants performed this task above chance on all three days ($p < 0.001$ for all days). A repeated measures ANOVA revealed that performance differed between days ($F_{2,44} = 5.24$, $p = 0.009$). This difference is driven by a difference in performance between days 1 and 3 ($p = 0.009$, post hoc tests using Bonferroni correction).

### Supplementary Figure 2

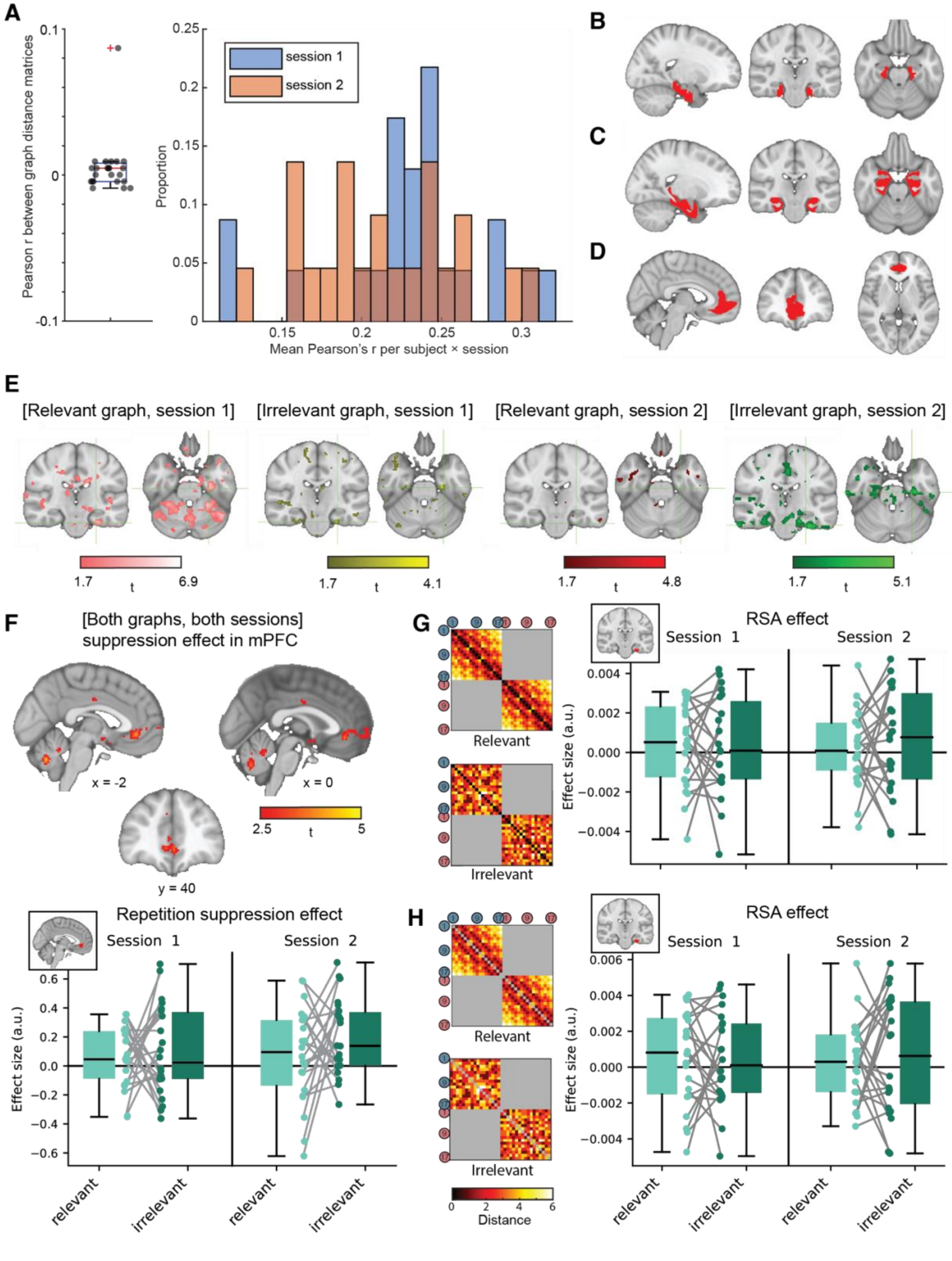

**Figure S2. Supplementary figures relating to the [both graphs, both sessions] repetition suppression contrast (related to Figure 3). A.** The distances on the “relevant” and “irrelevant” graphs were orthogonalised by design. Left: correlation between the distance matrices from each graph (each point is a participant). Right: Histogram of correlations between the HRF-convolved regressors (for each combination of subject, session and run) parametric regressors of distances on the two graphs. **B-D:** Anatomically defined regions of interest used for small-volume correction. **B.** Mask comprising bilateral entorhinal cortex and subiculum, received with thanks from Martin Chadwick (Chadwick et al., 2015). **C.** Mask comprising bilateral entorhinal cortex, hippocampus and parahippocampal cortex. Regions were defined using the maximum probability tissue labels provided by Neuromorphometrics, Inc (http://Neuromorphometrics.com). **D.** Medial prefrontal cortex mask, obtained by combining the masks of Brodmann areas 32sg and 14m from the Brainnetome atlas (Fan et al., 2016), thresholded at 50. **E.** Effects of individual conditions for the 2x2 design of graph relevance x session, aligned anatomically to the peak of the [both graphs, both sessions] MTL effect ([-24, -22, -28], displayed at $p < 0.05$ uncorrected for visualisation) . All individual effects, and pairwise differences between them, were not significant when corrected for multiple comparisons in either of the anatomical MTL masks used in the manuscript. **F.** [Both graphs, both sessions] effect in the mPFC. Top: Brain regions representing relevant and irrelevant distances across both sessions, displayed at $p < 0.01$ uncorrected for visualization. The cluster in the mPFC trended towards significance ($p = 0.055$ corrected at the cluster level in the mPFC mask depicted in **D**, peak $t_{22} = 4.13$ [-2, 38, -6]). Bottom: Visualisation of parameter estimates extracted from the mPFC cluster (inset). A 2x2 ANOVA with the factors “Graph relevance” and “Session” did not reveal any significant main effects or interactions (Session: $F_{1,21} = 1.99$, $p = 0.17$; Relevance: $F_{1,21} = 1.37$, $p = 0.26$; Session × Relevance: $F_{1,21} = 0.36$, $p = 0.55$). **G-H.** [both graphs, both sessions] RSA analysis. **G.** Left: Model RDMs (see Figure 4F in the main text for a schematic illustration of building the Data RDM): relevant and irrelevant distances, only comparing conditions from the same graph (grey elements are ignored). Right: Visualisation of parameter estimates for the RSA analysis regressing the data RDM from the left MTL repetition suppression cluster (inset) in Figure 3B onto the relevant and irrelevant model RDMs. As in the repetition suppression lMTL effect, a 2x2 ANOVA with the factors “Graph relevance” and “Session” did not reveal any significant main effects or interactions (Session: $F_{1,21} = 0.28$, $p = 0.6$; Relevance: $F_{1,21}1 = 0.001$, $p = 0.96$, Session × Relevance: $F_{1,21} = 1.5$, $p = 0.7$**). H.** Same as in G, but with the (visually confounded) diagonal ignored (NaNed out). Boxplots in F, G, H show the median, interquartile range, and individual participant values for each combination of Session and Graph relevance.

### Supplementary Figure 3

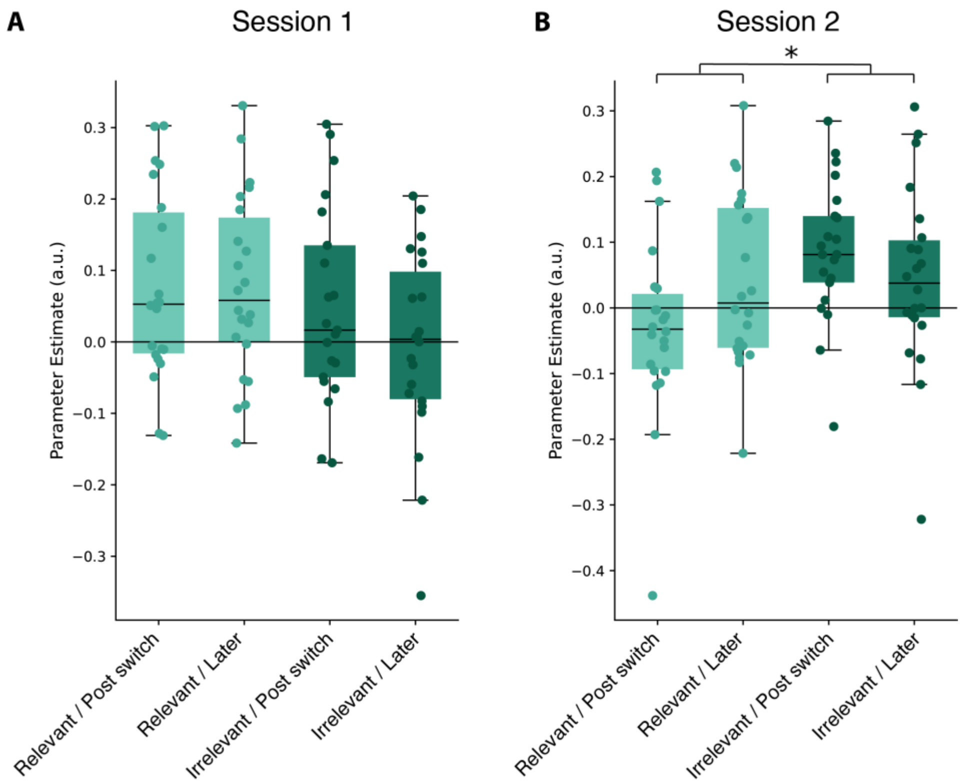


**Figure S3. Exploratory analysis of switch dynamics in hippocampus.** To examine whether relational coding differed as a function of trial position relative to a context switch, we conducted an exploratory 2 × 2 repeated-measures ANOVA (Relevance: relevant vs. irrelevant distance × Switch: first trial after switch vs. subsequent trials) separately for each session within the hippocampal ROI. **A** No significant interaction was observed in session 1 ($F_{1,21}$ = 1.74, p = 0.20). **B** In session 2, this analysis revealed a significant Relevance × Switch interaction ($F_{1,21}$ = 5.12, p = 0.034), indicating that the effect of relational relevance depended on trial position relative to the switch. Specifically, immediately following a switch, irrelevant distance coding was relatively enhanced while relevant distance coding was attenuated, whereas this difference was not present on subsequent trials.
